# Genes associated with fitness and disease severity in the pan-genome of mastitis-associated *Escherichia coli*

**DOI:** 10.1101/2023.07.26.549771

**Authors:** Michael A. Olson, Caz Cullimore, Weston D. Hutchison, Aleksander Grimsrud, Diego Nobrega, Jeroen De Buck, Herman W. Barkema, Eric Wilson, Brett E. Pickett, David L. Erickson

## Abstract

Bovine mastitis caused by *Escherichia coli* may manifest as subclinical through severe acute disease and can be transient or persistent in nature. Little is known about bacterial factors that impact clinical outcomes or allow some strains to outcompete others in the mammary gland (MG) environment. Mastitis-associated *E. coli* (MAEC) may have distinctive characteristics which may contribute to the varied nature of the disease. In this study, we sequenced the genomes of 96 MAEC strains isolated from cattle with clinical mastitis (CM). We utilized clinical severity data to perform genome-wide association studies to identify accessory genes associated with strains isolated from mild or severe CM, or with high or low competitive fitness during *in vivo* competition assays. Genes associated with pathogenic or commensal strains isolated from bovine and avian sources were also identified. A type-2 secretion system (T2SS) and a chitinase (ChiA) exported by this system were strongly associated with pathogenic isolates compared with commensal strains. Strains carrying these genes also had higher competitive fitness during experimental intramammary infections. Deletion of *chiA* from MAEC isolates decreased their adherence to cultured bovine mammary epithelial cells, suggesting that the increased fitness associated with strains possessing this gene may be due to better attachment in the MG.

**Importance:** Bovine mastitis caused by MAEC compromises animal health and inflicts substantial product losses in dairy farming. Given their high levels of intraspecies genetic variability, virulence factors of commonly used MAEC model strains may not be relevant to all members of this group. Here we analyzed clinical data as well as fitness (quantified in a mouse MG model) of diverse MAEC isolates to identify accessory genes that contribute to infection. We demonstrated a novel role for chitinase in promoting attachment to mammary epithelial cells. Reverse genetic approaches can be applied to the collection of strains and their complete genome sequences that we have presented here. Overall, these results provide a much richer understanding of MAEC and suggest bacterial processes that may underlie the clinical diversity associated with mastitis and their adaptation to this unique environment.

## Introduction

Bovine mastitis often occurs as a result of bacterial infection. Mastitis-associated *E. coli* (MAEC), abundant in the dairy environment, are the most important cause of this disease. These bacteria cycle between the bovine digestive tract and soils and beddings of stalls, from which they may gain access to the mammary gland (MG) via the teat canal. Once established within the MG, MAEC can induce a range of clinical presentations. Subclinical mastitis is usually defined as increased somatic cell counts in milk and transient inflammation caused by cytokine release. MAEC infections are typically cleared rapidly without complication or need for antibiotic intervention (1, 2). Conversely, severe clinical mastitis (CM) can damage the MG through sustained inflammation and high bacterial loads. These cows often suffer permanent udder damage, and the bacteria occasionally disseminate beyond the MG leading to sepsis (3, 4). Some mastitis cases are characterized by mild acute disease followed by extended periods of chronic or recurrent infections. Occasionally MAEC strains gain access through the teat canal to the MG during the non-lactating period. Symptoms of CM develop shortly after the next lactation begins, which also tends toward chronic infections (5, 6).

Features that distinguish MAEC from other *E. coli* strains have been difficult to identify and remain incompletely understood. Although they may come from any of the diverse *E. coli* lineages, MAEC strains often fall within *E. coli* phylogroups A and B1, which are also the most common phylogroups of commensal strains. Nevertheless, commensal strains are unable to cause acute clinical or chronic mastitis (7), suggesting that there are fundamental differences between commensals and MAEC strains that have yet to be discovered. MAEC frequently belong to sequence types (MLST) 10, 58, 95, and 1125 (8–13) but this also does not distinguish them from other strains. Genes predicted to be associated with MAEC have been examined in numerous PCR-based surveys as well as more extensive genomic studies (7, 11, 14–16).

Phenotypes distinctive of MAEC strains include relatively robust resistance to the complement system and greater motility than other *E. coli* (16–18). The ferric dicitrate transport system encoded by the *fecABCDE* genes is also highly expressed and much more consistently found in MAEC genomes compared with other *E. coli* (19, 20). Previously, we conducted a functional genetic screen using transposon insertion sequencing to identify the fitness factors of a single MAEC strain (21). This work demonstrated that the *fec* genes are needed to colonize lactating mouse MGs and implicated the high-affinity zinc transport system and several genes involved in other metabolic pathways in fitness in MGs. However, different genes may be required for fitness in other MAEC strain backgrounds. A more thorough understanding of MAEC genomics may help identify strains capable of causing mastitis from the varied strains found in agricultural settings, as well as those more likely to cause severe CM.

Mastitis severity depends on several host factors that contribute to disease outcome. For instance, the stage of lactation when the MG becomes infected has a strong influence on whether the infecting bacteria are efficiently cleared. At the time of parturition and earlier stages of lactation, cows are more prone to severe CM, which has been attributed to immunological dysfunction of neutrophils and lymphocytes during this time (22–27).

In addition to host factors, bacterial factors may also influence CM severity (17, 28, 29). Some MAEC strains carry virulence factors often found in human or avian extraintestinal pathogenic *E. coli* (ExPEC) such as toxins, siderophores, capsules and adhesins (30). For the most part, the influence of these virulence factors during intramammary infections has not been determined. Thus far, the only trait with a demonstrated association with CM severity is swarming motility, which is higher in MAEC strains isolated from severe CM cases than those strains isolated from mild and moderate CM (17). Gene expression comparisons of MAEC isolates from transient infections and persistent CM also demonstrated that flagella gene expression and motility are generally higher in the persistent CM strains (16). Persistent strains are also more resistant to serum complement and express the *fec* operon genes at higher levels than transient strains.

Previous genomic analyses have focused on identifying genes that are unique to MAEC isolates compared to non-pathogenic strains inhabiting the same niches or commensal strains belonging to the same phylogroups. These analyses have uncovered putative marker genes that could distinguish MAEC from other strains, including those that may function in niche-specific metabolic pathways, gene regulation, and virulence (19, 31). In this study, we sequenced 96 MAEC genomes and implemented a comparative genomics approach to uncover genes more likely to be present in strains isolated from cattle with mild or severe CM. We then extended this analysis to identify genes associated with either pathogenic or commensal *E. coli* strains previously isolated from bovine and avian hosts. We employed mouse infection and milk growth assays to separate MAEC strains with higher or lower fitness to identify genes that are associated with these phenotypes.

## Materials and methods

### Bacterial strains and growth conditions

*E. coli* strains M22 through M117 (Supplementary Dataset 1) were isolated from individual quarter milk samples from cattle with clinical mastitis as part of the Canadian National Cohort of Dairy Farms (32). In brief, 89 herds across Canada (Alberta, Ontario, Quebec, and the Maritime provinces Prince Edward Island, New Brunswick, and Nova Scotia) were selected to be representative of their respective province in terms of housing type, bulk tank somatic cell count, cattle breed, and milking schedule, and were followed from February 2007 to December 2008. At the time of collection, farmers evaluated the clinical signs presented in each affected animal and assigned a clinical score, as follows: mastitis score 1 (mild = abnormal milk only), mastitis score 2 (moderate = abnormal milk and local inflammation signs), or mastitis score 3 (severe = abnormal milk, local inflammation and systemic clinical signs (33). From this collection (34), strains isolated from cows in the middle to late stages of their lactation cycle were selected in order to focus on bacterial differences and minimize the effect that early lactation has on mastitis severity (35). Selection was also designed to include isolates from different herds and provinces. Six strains isolated from CM cases in the United States were also included in this study (Supplementary Dataset 1). Strains M3, M6, M9, M11, and M12 were previously isolated from quarter milk samples of CM cases (21). Clinical severity data were not available for these isolates. Strain G1 was supplied by Jennifer Wilson (Jersey Girls, Jerome ID) and was isolated from a cow with severe, gangrenous mastitis that necessitated culling of the animal.

Putative *E. coli* were identified phenotypically by colony morphology on blood and MacConkey agar plates and confirmed by sequencing. Bacteria were routinely grown in Luria-Bertani (LB) medium at 37°C. For milk cultures, whole, unpasteurized cow’s milk was obtained from a local supplier and used immediately or stored at −80°C until use. To determine growth yields of individual MAEC isolates, bacteria from overnight LB cultures were added to 100 μl milk to a concentration of 10^3^ CFU/ml in a 96-well format. Plates were incubated without shaking at 37°C. A sample was immediately removed (T=0), serially diluted and plated on MacConkey agar to determine the starting concentration. Bacterial concentrations were also measured at 4 h and 8 h post-inoculation. The change in CFU/ml at 4 h and 8 h relative to T=0 was calculated for three biological replicates for each strain.

### Genome sequencing, assembly, and annotation

Total DNA was isolated from MAEC strains using a ZR Fungal/Bacterial DNA MiniPrep kit (Zymoresearch). DNA sequencing libraries were prepared using the Illumina Nextera DNA Library Prep kit as previously described (36). DNA libraries were sequenced by Genewiz, Inc. (South 330Plainfield, NJ), and Illumina paired-end reads of 150 bp were generated on a MiSeq with version 2 chemistry. Contig assembly, *in silico* determination of phylogroup, multi-locus sequence typing, and GrapeTree analysis were performed using EnteroBase (37, 38) and genomes were annotated using Prokka (version 1.14.6) (39).

### Core and accessory genome determination, alignment, phylogenetic trees, and pan-genome analysis

PIRATE (version 1.0.4 with default parameters) was used to perform a pan-genome analysis on all the GFF annotation files for all MAEC genomes and the results were then converted to the ROARY gene presence/absence format. The core genome alignment was used to create a maximum-likelihood phylogenetic tree using IQ-Tree, and the subsequent tree was visualized using Interactive Tree of Life (40–42). SCOARY (version 1.6.16 with default parameters) was then run on the converted PIRATE output to identify genes found in the accessory genome. For the severity analysis, we selected 81 isolates belonging to mastitis scores 1 or 3 (40 mild, 41 severe). SCOARY was also used to identify genes in the pan-genome associated with competitive fitness in milk and in mouse MGs (using barcoded strains, see below). For this analysis, the top and bottom 30% of strains for each condition were separated based on their CI values (regardless of whether they came from mild or severe CM cases). To identify genes associated with pathogenic or commensal strains, genomes for bovine commensal strains were downloaded from NCBI using “bos taurus commensal” with the *E. coli* species tag. Similarly, previously uploaded MAEC genomes were downloaded from NCBI using “bovine mastitis” with the *E. coli* species tag or from (43) using the python downloading programs at https://github.com/SomeoneNamedCaz/E.-Coli-genome-analysis. Genome sequences for avian strains (pathogens and commensals) were obtained from Mageiros et al (44). SCOARY input files consisted of a gene absence/presence Rtab file generated by PIRATE and a custom trait file which assigned a discrete phenotype for each strain.

### Hierarchical clustering analysis

The MD Anderson Cancer Center Next-Generation Clustered Heat Map (NG-CHM) builder was used for hierarchical clustering (https://build.ngchm.net/NGCHM-web-builder/) based on the ExPEC virulence gene carriage. The Euclidian distance and single-linkage methods were used to generate the clusters.

### Detection of plasmids and antimicrobial resistance (AMR) genes

The PlasmidFinder 2.1 database at the Center for Genomic Epidemiology (https://cge.cbs.dut.dk/services/PlasmidFinder) was used to detect and type plasmids found in MAEC genomes. Assembled reads for each strain were searched using the most recent Enterobacteriaceae plasmid database using 90% minimum identity and 60% minimum % coverage cutoffs. Incompatibility (Inc) groups that were found in each strain were recorded. Each Inc group was counted individually even when multiple Inc groups were detected in a single strain. ABRicate software (https://github.com/tseemann/abricate) was used to find genes implicated in AMR from the Resfinder database (45).

### Development of barcoded plasmids and barcode sequencing

The low copy pACYC184 plasmid was modified by designing PCR primers incorporating a partial Illumina adapter flanking 12 random nucleotides and eliminating the tetracycline resistance gene, yielding a 2102 bp product (Supplementary Figure 1). The left primer incorporated the random sequences and the first Illumina Truseq adapter, and both primers included SalI overhang sequences. The PCR product was digested with SalI, ligated and transformed into competent *E. coli* DH5α. Ninety-six unique plasmids were isolated, and their barcode sequence determined by Sanger sequencing. These plasmids were then transformed into individual MAEC isolates. Four strains were unable to be transformed because of pre-existing chloramphenicol resistance, leaving 92 strains that were successfully transformed. An inoculum was prepared by growing each strain individually in LB to an A_600_=1.0, mixing 20 μl of each strain, together, and diluting the mixture to a final concentration of 5×10^6^ CFU/ml in PBS. This inoculum was frozen for use in competition tests and was sequenced as the input library.

The input library was grown in duplicate in LB broth and in whole unpasteurized cow milk (pooled from healthy cows from a local commercial supplier) for 8 h. The input library was also injected into lactating mouse MGs (see below). After growth in LB, milk and MG infections, plasmids were isolated from the bacterial population in each sample. The barcodes were amplified using primers that also added the remainder of the Illumina adapter as well as sample-specific identifying sequence. Illumina paired-end reads of 150 bp were generated on MiSeq version 2 sequencer by Genewiz, Inc. (South Plainfield, NJ) and CD-Genomics, Inc. (Shirly, NY). A custom grep function was used to identify and count barcodes for each strain from the sequence reads. Fitness scores were calculated as the number of reads for each barcode in a output sample as a proportion of the total reads (all barcodes) in that sample, divided by the ratio of that same barcode to the total reads in the inoculum (input) library.

### Mouse infections

Lactating CD-1 IGS mice between 9 and 12 weeks of age and 10 to 11 days postpartum were infected as previously described (21). Briefly, a 50 µl volume of bacteria containing 500 CFU of each strain for total of ∼50,000 CFU was suspended in PBS. The inoculum was injected directly through the teat canal into the ductal network of the 4^th^ left and 4^th^ right MGs of individual mice using a 33-gauge needle with a beveled end. Pups were removed for 1 to 2 h after injections and then reunited with the mother and allowed to nurse normally. Mice were euthanized 24 h post infection and MG tissue was homogenized in 1 mL PBS. The tissue homogenate was added to LB broth containing chloramphenicol (10 μg/ml) to recover the bacteria and isolate plasmid DNA.

### Deletion and complementation of *chiA*

An allelic exchange plasmid was created using the pAX1 plasmid (46). Upstream and downstream regions (500 bp) of *chiA* were amplified, stitched together using overlap extension, and inserted into the SalI and AvrII sites of the pAX1 plasmid. The resulting plasmid was transformed via electroporation into the donor *E. coli* strain MFDλpir. Integration of the suicide plasmid to create merodiploid recipients and excision to produce unmarked deletions of chiA in each of the strains was performed as described (46). Complementation of *chiA* gene was done by amplifying *chiA* including 300 bp upstream to include the putative promoter and ligating the resulting product into pJET1.2. The resulting plasmid pWH01 was verified by sequencing and transformed via electroporation into each deletion mutant.

### MAC-T Cell Culture and Media

Bovine mammary alveolar epithelial cells (MAC-T cells) were generously provided by Dr. Janos Zempleni (University of Nebraska-Lincoln). They were grown in T-75 flasks with 40% (v/v) Dulbecco’s Modified Eagle Medium (DMEM), 40% (v/v) Ham’s F12 Medium, and 10% (v/v) bovine serum (FBS). FBS was heat-inactivated prior to use in media by incubating in a 56°C water bath for 30 minutes with periodical mixing. This was supplemented with 5 μg/mL bovine insulin, 1 μg/mL hydrocortisone, 23 mM HEPES buffer, 2.2 g/L sodium bicarbonate, and 40 mM L-glutamine. Penicillin, streptomycin (100 U/mL and 100 μg/mL, respectively) and amphotericin B (2.5 μg/mL) were added into the media for routine growth. MAC-T cells were grown at 37°C and 5% CO_2_ (v/v).

### Adhesion assays

MAC-T cells were seeded into a 12-well plate and grown to ≥95% confluency. The density of epithelial cells was determined by trypan blue exclusion and counting using Cell Counter model R1 automated cell counter (Olympus). Approximately 6×10^5^ cells were present in each well. The number of cells and viability did not vary throughout the assays as determined by trypan blue exclusion. On the day of each assay, spent media was removed from each well and the cells were washed 3 times with 1 mL of sterile PBS to remove residual antibiotics from media. Media without antibiotics was added into each well and cells were allowed to incubate for ≥2 hours prior to inoculation with bacteria.

Overnight cultures of bacteria were diluted in sterile PBS to an OD_600_ of 0.5. Bacteria were added to a multiplicity of infection (MOI) of 10:1. The 12-well plates were then centrifuged at 300 x g for 5 min at room temperature to synchronize contact of bacteria with the MAC-T cells. Plates were then incubated at 37°C and 5% CO_2_ (v/v) for 1 h. Following incubation, media was aspirated and wells washed 3× with 1 mL of sterile PBS to dislodge non-and weakly adhered bacteria. Triton X-100 (500 μL, 0.1%) in PBS was then added into each well and incubated at room temperature for 5 minutes to lyse cells. The resulting suspension of bacteria was then homogenized, serially diluted and plated on LB agar overnight and CFUs were counted.

### Statistical analyses

Statistical analyses were carried out using Prism9 (GraphPad) or SCOARY. A *P*-value <0.05 was considered statistically significant. For GWAS studies, Fisher’s exact test was used to determine significance for each gene. Associations of the Inc group with a strain type (mild or severe CM) were assessed using Fisher’s exact test. Distribution of AMR genes between mild and severe CM isolates was compared using the Mann-Whitney T-test. Correlations between fitness scores were determined by Spearman rank correlation analysis or Mann-Whitney T-test. Differences in adherence to MAC-T cells by wild-type, *chiA* mutant, or complemented strains were analyzed using a one-way analysis of variance (ANOVA).

## Results

### Bacterial growth in milk is not associated with CM severity

The ability to efficiently utilize the nutritional components present in milk and thus grow rapidly may be a determining factor for some strains ability to colonize MGs successfully. To determine if there is a link between *in vitro* growth in milk and CM severity, we compared the growth of a subset of mild and severe MAEC strains in whole unpasteurized cow’s milk (Supplementary Figure 2). Replication was measured after four or eight hours of growth. MAEC strains varied widely in their growth yields, with some strains replicating 100-fold within four hours and up to 100,000-fold by eight hours, whereas others did not increase. However, replication was not different between strains isolated from mild and severe CM at either time point. This is consistent with other reports that *in vitro* growth in whole milk is not associated with CM severity (47).

### Genome analysis of mild and severe CM isolates

Complete genome sequences for 81 MAEC strains (41 severe and 40 mild CM isolates) were assembled and annotated (Supplementary Dataset 1). From these annotated sequences, we identified a total of 15,395 unique genes representing the pan-genome (48). Of these, 3,177 (20.6%) were considered core genes (>98% prevalence), 263 (1.7%) were considered soft core genes (95-98% prevalence), 1,523 were considered shell genes (15-95% prevalence), and 10,432 (67.8%) were considered cloud genes (<15% prevalence) (Supplementary Table 1).

*In silico* phylogroup analysis demonstrated that most strains belong to phylogroups A and B1 (49). Multi-locus sequence typing based on seven house-keeping genes was also used to more precisely assess the phylogenetic backgrounds of the MAEC strains (50). A total of 30 different STs were identified, with many STs being represented by only one strain each. In our study, ST10, ST58, ST1121 and ST1125 were the most abundant, representing 45% of the total strains (Figure 1a). MLST groupings did not correlate with mild or severe CM. In previous genomic studies of MAEC strains, most were classified as ST10, ST23, ST58, ST88, or ST1125 in phylogroups A or B1, which is consistent with our findings (9–11).

**Figure 1.**
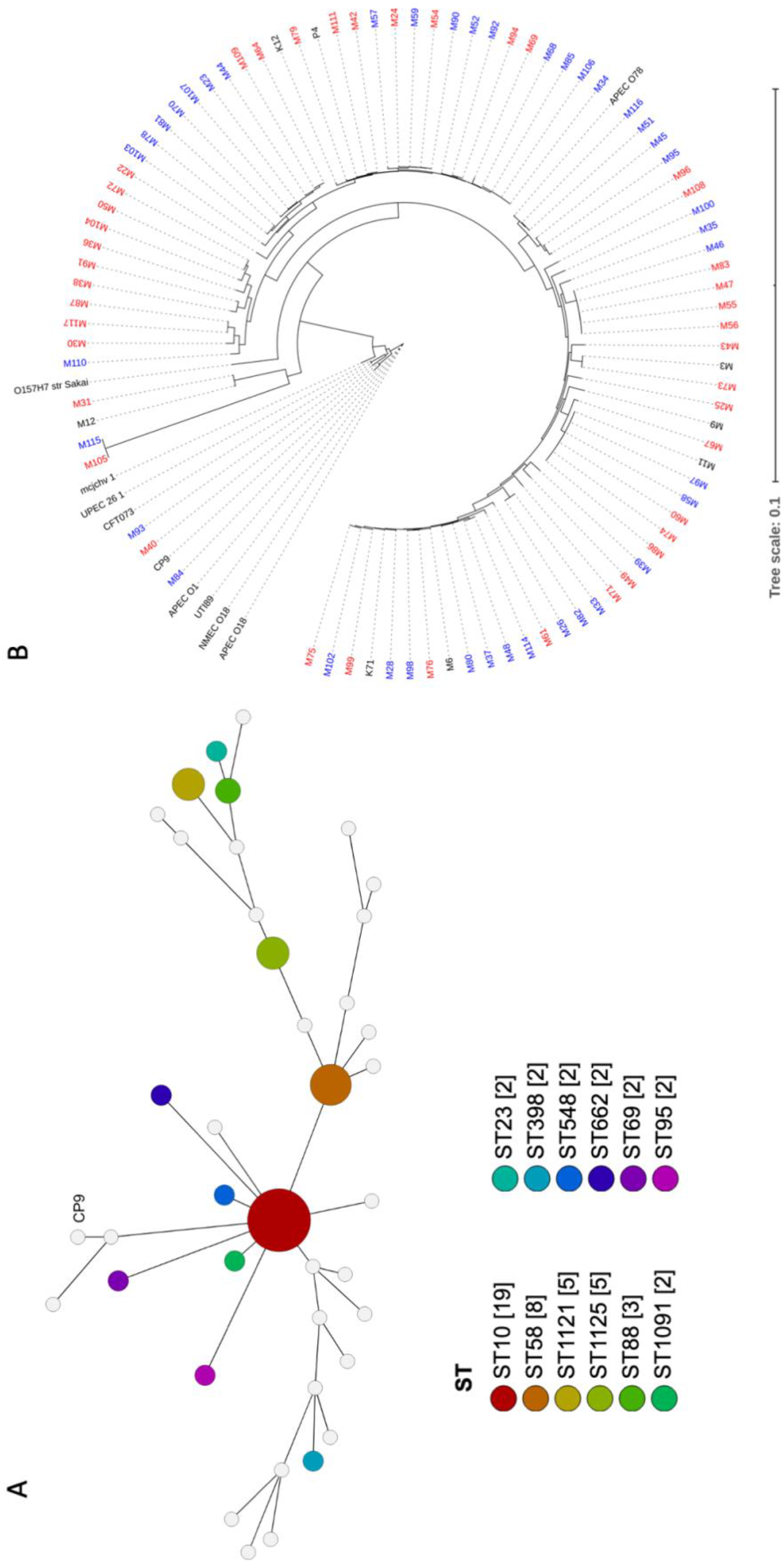
Genome comparison of MAEC strains. (A) Sequence type (ST) prediction using the GrapeTree function with the MSTree V2 algorithm to visualize strain relatedness (37, 38). Overall, 30 unique STs were detected. The figure displays STs represented more than once in the population (node diameter scaled to frequency of detection). Human ExPEC strain CP9 was also included for comparison. (B) Core genome relatedness among strains isolated from severe mastitis or mild mastitis. Strains isolated from mild CM cases are denoted in blue and severe CM in red. Strains labeled black were included as references or did not have clinical severity data and were not part of the analysis.

To explore whether CM severity can be predominantly attributed to differences in the core genome, a phylogenetic tree was constructed based on the core genome sequence alignments. In this analysis, several ExPEC strain, one enterohemorrhagic strain, and additional MAEC strains without clinical severity data were included as references (Figure 1b). The phylogenetic tree reveals the high diversity of MAEC strains. Compared to the relatively tight clustering of other ExPEC, the MAEC strains belong to a broad range of backgrounds, including some closely related to other ExPEC strains. Of 10 strains that clustered closely with other ExPEC strains, nine were isolated from severe CM and one from mild CM. However, there was no consistent clustering into clades based on CM severity. There was also no association between the core genome and the geographical region where the strains were isolated (Supplementary Figure 3).

ExPEC are difficult to define based on any single or group of virulence genes. However, they often carry genes related to iron acquisition (yersiniabactin *ybtP*, aerobactin *iutA*, salmochelin *iroN* siderophores, ferric citrate *fecA* and *sit* ferrous iron transport), group 2 or 3 capsules (*kps*), adhesive and invasive factors (P fimbriae; *papC,* S fimbriae; *sfaA*, *focC*, afimbiral adhesins (*afaD*), and toxins (*sat*, *hlyA*, *cdtA*) (51–55). These genes were detected in several of the MAEC genomes (Figure 2). Mean number of virulence genes carried by each strain was 3.32±2.15 or 3.17±2.23 for mild and severe isolates, respectively, demonstrating that simple abundance of ExPEC virulence genes is not predictive of CM severity. For example, strain M93 (mild isolate) carries 10 of these ExPEC virulence genes. As CM severity could be associated with specific combinations of virulence genes, hierarchical clustering was then performed based on the presence or absence of each gene (Figure 2). This analysis demonstrated no apparent clustering of strains based on CM severity and ExPEC virulence gene content.

**Figure 2.**
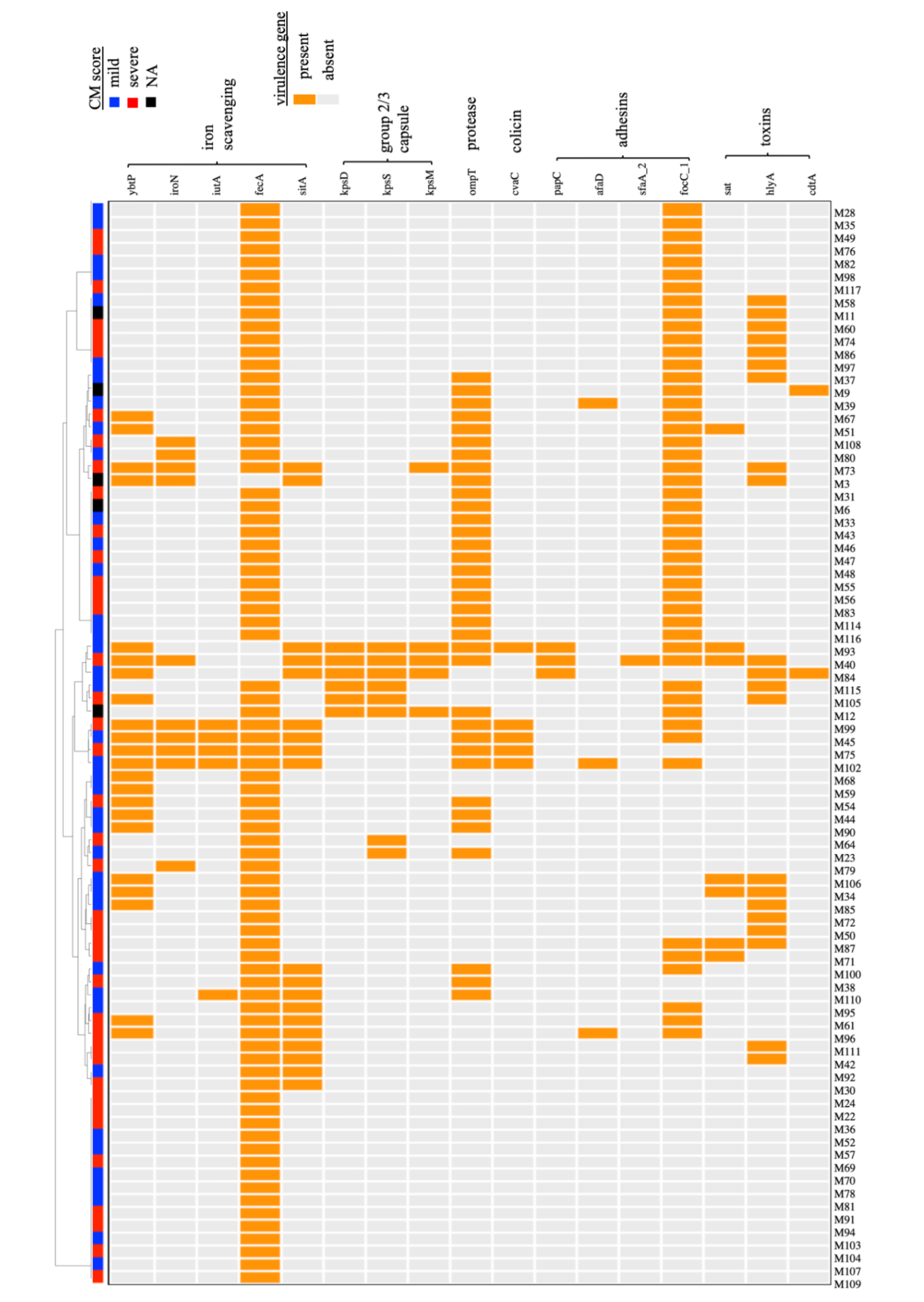
Hierarchical clustering of MAEC strains based on the carriage of virulence genes associated with the ExPEC phenotype. A dendogram was built based on the presence or absence of each gene. Both mild and severe CM isolates were present in each clade, and no relationship between CM severity and particular combinations of virulence genes was detected. NA-CM severity score not available.

Plasmids contribute to bacterial diversity such as that demonstrated by our study population, in part because they can contribute to homologous or non-homologous recombination. Plasmids also frequently carry specific fitness or AMR genes, which could influence CM severity. Incompatibility typing was performed for each of the MAEC strains (56, 57). Nineteen different incompatibility (Inc) groups were detected in the MAEC genome sequences, suggesting a wide diversity of plasmid content. Multiple Inc types were frequently detected in the same strain, suggesting the possibility of hybrid plasmids. When compared to the severe CM strains, more Inc groups were detected among the mild CM strains (Figure 3). IncF1B was the most detected group in both mild and severe CM strains. IncF1B and IncF1C were more abundant in mild strains than in severe CM strains.

**Figure 3.**
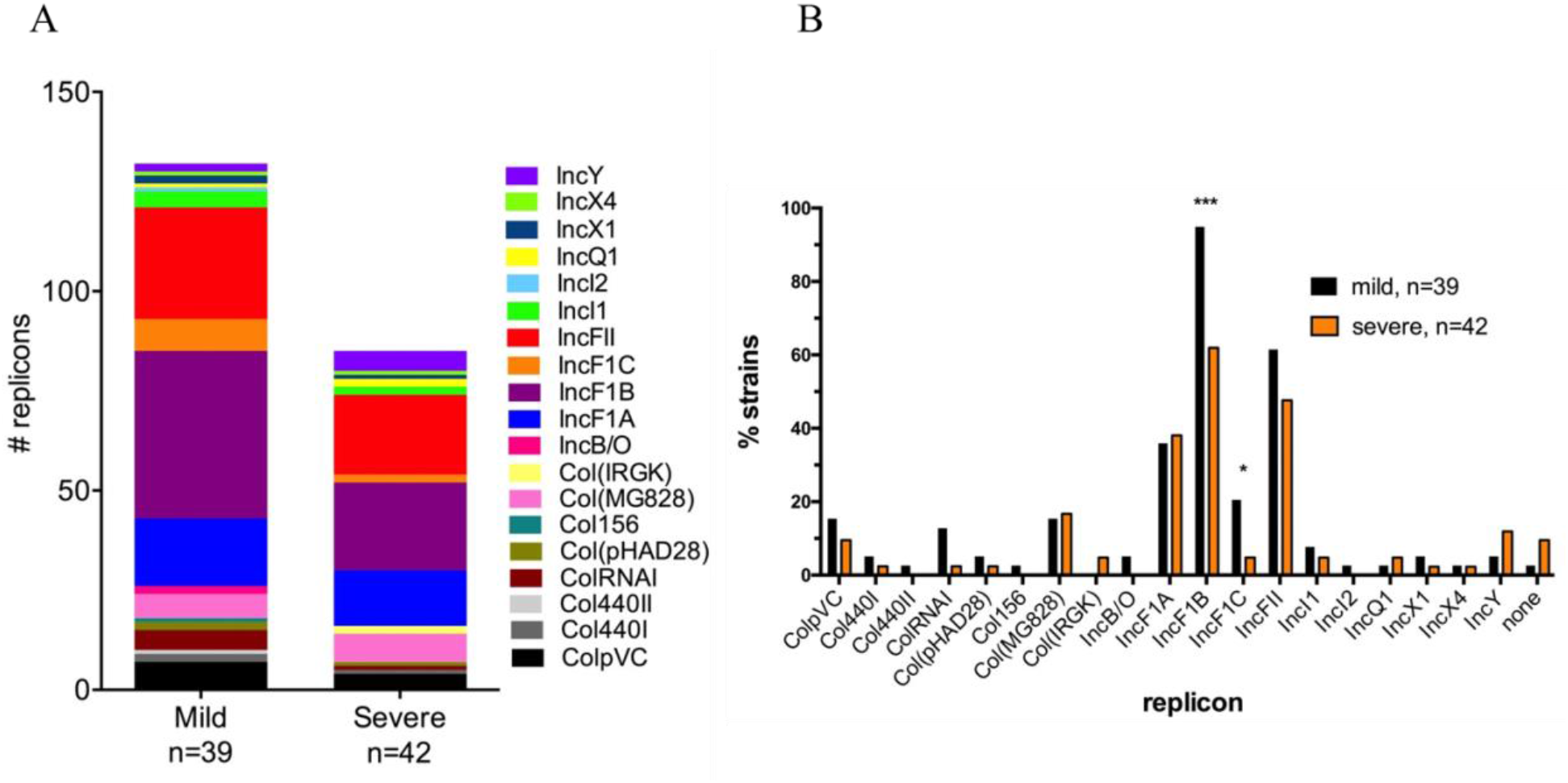
Plasmid replicons (Inc groups) found in mild and severe CM strains. (A) Total plasmid replicon genes detected in mild and severe strains. More unique replication genes were detected in mild CM strains overall compared with severe CM strains. (B) The proportion of strains that carry each type of replicon. Mild strains were significantly more likely to possess IncF1A and IncF1C replicons when compared to severe strains (\*\*\**P*<0.001, * *P*<0.05 by Fisher’s exact test).

Carriage of AMR genes among mild and severe CM strains was also assessed *in silico* by searching a repository of AMR genes (58). This analysis demonstrated that 12 strains carried one or more genes conferring resistance to aminoglycosides, beta-lactams, anti-folates, macrolides, phenicols, and tetracyclines (Supplementary Table 2). Aminoglycoside and anti-folate resistance genes were the most abundant. Notably, two severe CM isolates (M79 and M96) had nine AMR genes each. However, the distribution of AMR genes was not significantly different between mild and severe CM isolates (*P* =0.6029).

### Genes associated with CM severity

Next, we analyzed each gene in the pan-genome and scored it according to the association with mild or severe CM phenotypes. Each gene with an apparent association with the phenotype was reanalyzed, incorporating information about the phylogenetic structure to implicate genes associated with CM severity (59). Using a *P*-value cutoff of 0.05, we detected 25 genes that were associated with severe CM and 79 genes associated with mild CM (Supplementary Dataset 1 and Table 1). Among those associated with severe CM were two genes likely involved in O-antigen or capsule biosynthesis (*wbpI* and *capD*) and several genes involved in producing Yad fimbriae. These genes were detected in eight strains isolated from severe CM cases and were not present in any strains from mild CM cases. Yad fimbriae are composed of the tip/adhesin YadC along with fimbrial protein subunits encoded by *yadMLKN* genes. Assembly requires the chaperone protein YadV and the fimbrial usher HtrE (60–62). Of these genes, *htrE, yadM, yadL, yadK, yadN,* and *yadV* were all positively associated with severe CM.

**Table 1.**
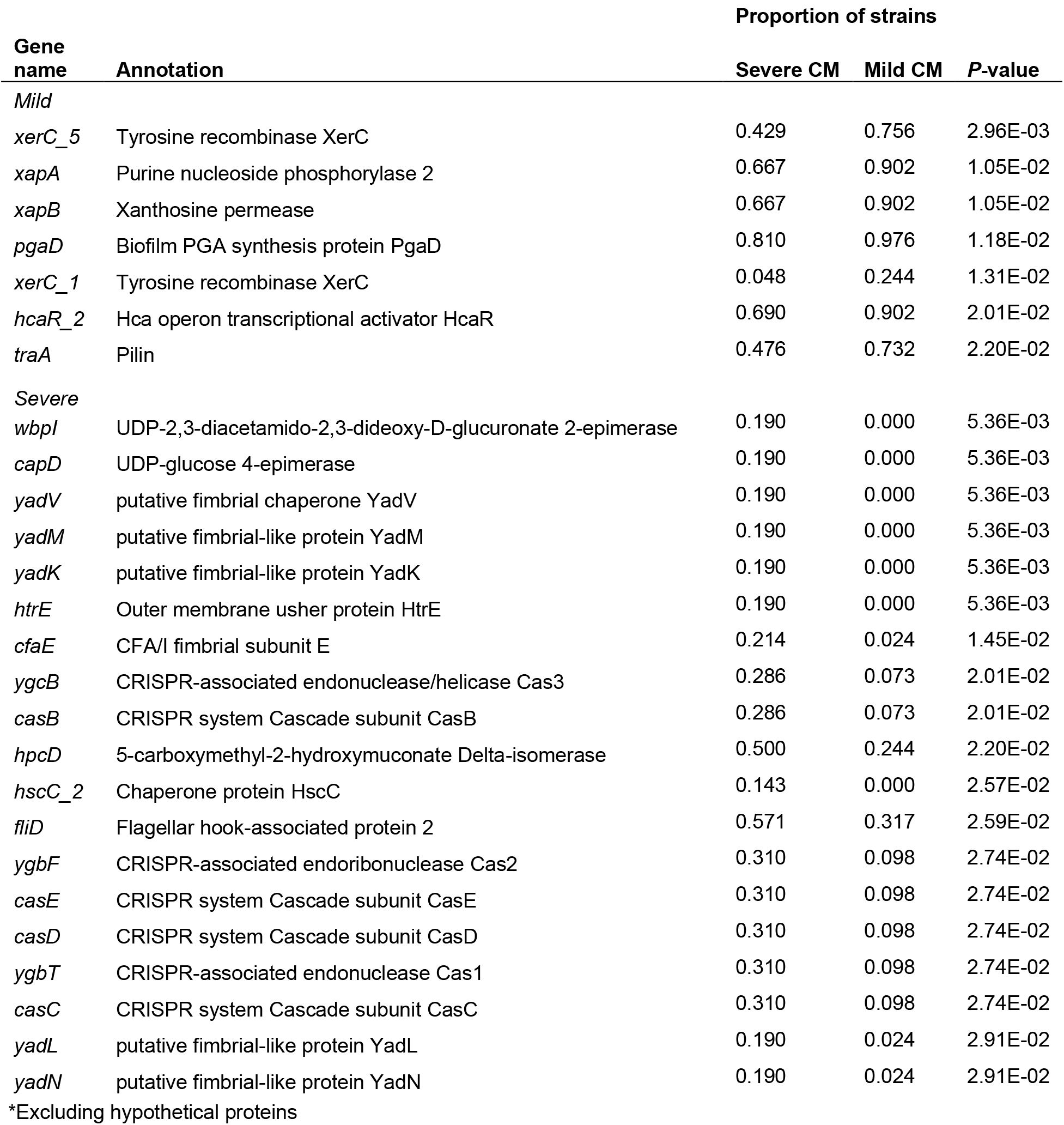
Top genes* associated with MAEC isolated from mild or severe clinical mastitis (CM)

| Gene name | Annotation | Proportion of strains |  |  |
| --- | --- | --- | --- | --- |
|  |  | Severe CM | Mild CM | P-value |
| Mild |  |  |  |  |
| *xerC_5* | Tyrosine recombinase XerC | 0.429 | 0.756 | 2.96E-03 |
| *xapA* | Purine nucleoside phosphorylase 2 | 0.667 | 0.902 | 1.05E-02 |
| *xapB* | Xanthosine permease | 0.667 | 0.902 | 1.05E-02 |
| *pgaD* | Biofilm PGA synthesis protein PgaD | 0.810 | 0.976 | 1.18E-02 |
| *xerC_1* | Tyrosine recombinase XerC | 0.048 | 0.244 | 1.31E-02 |
| *hcaR_2* | Hca operon transcriptional activator HcaR | 0.690 | 0.902 | 2.01E-02 |
| *traA* | Pilin | 0.476 | 0.732 | 2.20E-02 |
| Severe |  |  |  |  |
| *wbpl* | UDP-2,3-diacetamido-2,3-dideoxy-D-glucuronate 2-epimerase | 0.190 | 0.000 | 5.36E-03 |
| *capD* | UDP-glucose 4-epimerase | 0.190 | 0.000 | 5.36E-03 |
| *yadV* | putative fimbrial chaperone YadV | 0.190 | 0.000 | 5.36E-03 |
| *yadM* | putative fimbrial-like protein YadM | 0.190 | 0.000 | 5.36E-03 |
| *yadK* | putative fimbrial-like protein YadK | 0.190 | 0.000 | 5.36E-03 |
| *htrE* | Outer membrane usher protein HtrE | 0.190 | 0.000 | 5.36E-03 |
| *cfaE* | CFA/I fimbrial subunit E | 0.214 | 0.024 | 1.45E-02 |
| *ygcB* | CRISPR-associated endonuclease/helicase Cas3 | 0.286 | 0.073 | 2.01E-02 |
| *casB* | CRISPR system Cascade subunit CasB | 0.286 | 0.073 | 2.01E-02 |
| *hpcD* | 5-carboxymethyl-2-hydroxymuconate Delta-isomerase | 0.500 | 0.244 | 2.20E-02 |
| *hscC_2* | Chaperone protein HscC | 0.143 | 0.000 | 2.57E-02 |
| *fliD* | Flagellar hook-associated protein 2 | 0.571 | 0.317 | 2.59E-02 |
| *ygbF* | CRISPR-associated endoribonuclease Cas2 | 0.310 | 0.098 | 2.74E-02 |
| *casE* | CRISPR system Cascade subunit CasE | 0.310 | 0.098 | 2.74E-02 |
| *casD* | CRISPR system Cascade subunit CasD | 0.310 | 0.098 | 2.74E-02 |
| *ygbT* | CRISPR-associated endonuclease Cas1 | 0.310 | 0.098 | 2.74E-02 |
| *casC* | CRISPR system Cascade subunit CasC | 0.310 | 0.098 | 2.74E-02 |
| *yadL* | putative fimbrial-like protein YadL | 0.190 | 0.024 | 2.91E-02 |
| *yadN* | putative fimbrial-like protein YadN | 0.190 | 0.024 | 2.91E-02 |

\*Excluding hypothetical proteins

Most genes associated with mild CM were hypothetical genes. However, genes for producing the biofilm PGA exopolysaccharide (63) and several tyrosine recombinases (64–67) were associated with mild CM. The complete *pgaABCD* operon was detected in nearly all (39 of 40) strains with a mild CM score whereas just *pgaD* was missing in eight strains from severe CM. Three tyrosine recombinase alleles (10, 31, 34) were associated with mild CM strains, while distinct alleles (2, 18, and 26) were associated with severe CM strains (Table 1).

### Genes associated with pathogenic vs commensal and avian vs bovine strains

To increase the power of our study, we conducted a complementary analysis of a larger publicly available set of *E. coli* genomes to identify genes that may distinguish commensal from pathogenic strains. These included strains isolated from the GI tracts of cattle, which were designated as commensals. Likewise, sequences for all strains with an identifiable mastitis designation as well as strains from this study or published by Alawneh et al. (43) were designated as pathogens. In order to identify genes that could contribute to disease in MGs that may also be relevant in other hosts or tissues, we also utilized sequences from a large collection of avian strains that were recently published (44) and separated into pathogens and commensals. Genes with significantly different distribution across categories (avian vs. bovine, pathogen vs. commensal) were identified.

Several genes were strongly associated with one animal host (avian) vs another (bovine) regardless of pathogenicity (Supplementary Dataset 1 and Table 2). For example, genes that comprise the hydrogenase-4 operon (*hyf*) including the putative formate transporter (*focB*) were almost universally present in bovine-associated strains (98.5%), compared with only 34% of avian strains. Hydrogenase-4 is involved in producing H_2_ and CO_2_ during fermentation in alkaline conditions as occur in the bovine rumen. Conversely, avian strains were far more likely to possess genes typically found on APEC plasmids such as salmochelin production and receptor genes, the *sitABCD* iron/manganese transport system, and the antibiotic efflux pump *etsA*.

**Table 2.** Top genes* associated with avian or bovine *E. coli* strains.

| Gene name | Annotation | Proportion of strains |  |  |
| --- | --- | --- | --- | --- |
|  |  | avian | bovine | P-value |
| *Avian* |  |  |  |  |
| O2ColV47 | Cobalamin (vitamin B12) biosynthesis CobW-like | 0.785 | 0.033 | 2.89E-136 |
| O2ColV54 | Multidrug efflux ATP-binding/permease protein | 0.798 | 0.051 | 2.68E-131 |
| *iroD* | Salmochelin esterase | 0.791 | 0.051 | 8.38E-129 |
| *iroB* | salmochelin biosynthesis C-glycosyltransferase | 0.791 | 0.051 | 8.38E-129 |
| *iroN* | Ferric salmochelin receptor | 0.787 | 0.058 | 4.17E-124 |
| *sitD* | Iron/manganese transport protein | 0.878 | 0.144 | 9.75E-123 |
| *sitB* | Iron/manganese transport protein | 0.880 | 0.149 | 1.12E-121 |
| *sitC* | Iron/manganese transport protein | 0.874 | 0.144 | 4.11E-121 |
| APECO1_2632 | putative isochorismatase hydrolase | 0.712 | 0.020 | 1.02E-120 |
| APECO1_2638 | Cytosine permease | 0.674 | 0.018 | 1.39E-111 |
| *Bovine* |  |  |  |  |
| *sfmF* | putative fimbrial-like protein | 0.281 | 0.967 | 9.68E-116 |
| *hyfF* | hydrogenase-4 NAD(P)H-quinone oxidoreductase subunit | 0.317 | 0.985 | 1.94E-115 |
| *fimZ* | two-component system, NarL family, response regulator | 0.253 | 0.949 | 5.42E-115 |
| *adiY* | HTH-type transcriptional activator | 0.321 | 0.980 | 1.45E-111 |
| *hyfD* | hydrogenase-4 NAD(P)H-quinone oxidoreductase subunit 2 | 0.342 | 0.985 | 1.04E-108 |
| *hyfE* | hydrogenase-4 component E | 0.342 | 0.985 | 1.04E-108 |
| *hyfB* | hydrogenase-4 component B | 0.342 | 0.985 | 1.04E-108 |
| *hyfG* | hydrogenase-4 component G | 0.342 | 0.985 | 1.04E-108 |
| *focB* | putative formate transporter 2 | 0.342 | 0.985 | 1.04E-108 |
| *manR* | Transcriptional regulator | 0.698 | 0.972 | 3.18E-108 |

\*Excluding hypothetical proteins

Analysis of the dataset for genes associated with pathogens vs commensals (regardless of host) revealed that the ferric dicitrate receptor genes (*fec* operon) were the most strongly associated with pathogenicity. Genes encoding the accessory type II secretion system and the *chiA* gene were also strongly associated with disease-causing strains (Table 3).

**Table 3.**
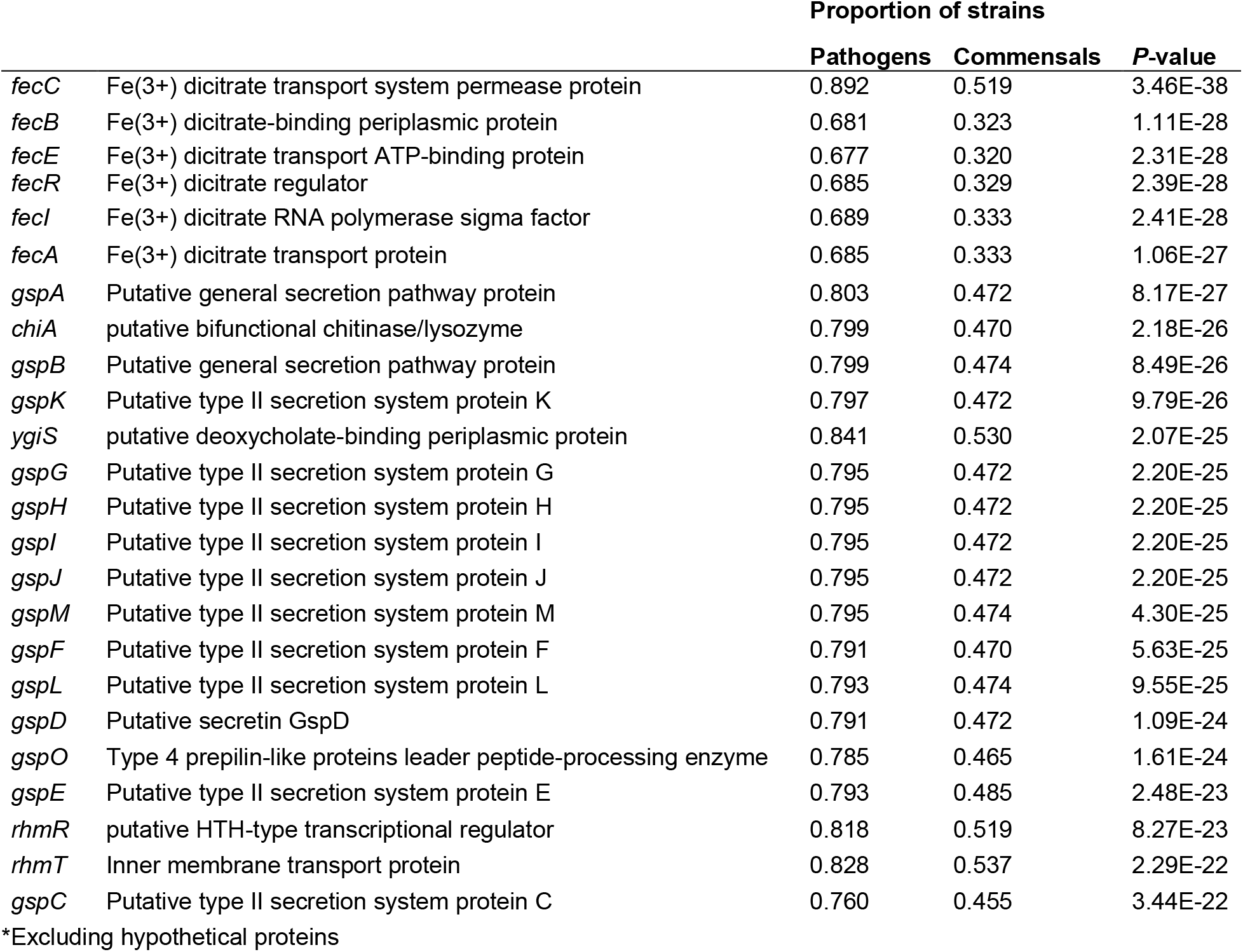
Top genes* associated with pathogenicity in bovine and avian-associated *E. coli*.

### Genes associated with fitness in milk and MGs

Next, we tested whether strains isolated from severe or mild CM have different fitness levels within the MG environment. We competed 92 MAEC strains against each other and used GWAS to identify genes associated with fitness. In these assays, pools of barcoded MAEC strains were grown in LB media, whole unpasteurized milk, and mouse MGs. Seventy-three bacterial strains were successfully recovered (sequencing failed for 19 barcodes and these strains were excluded from the analysis). From these 73 remaining strains, the abundance of each barcode was used to calculate the competition index (CI) as a measurement of fitness for each strain (Supplementary Dataset 1).

The CI scores that were calculated for each strain in duplicate LB and milk samples were highly consistent with each other (R^2^ = 0.99 and 0.94 respectively). The replicate samples for the nine MG infections were much more variable than in LB or milk. However, several strains consistently outcompeted others in mouse MGs. For instance, strain M65 exhibited a mean CI of 225. The mean CI of each strain during growth in LB and milk, (Figure 4a), milk and MGs (Figure 4b), and LB and MGs (Figure 4c) were positively correlated. Correlation between CI milk and MGs was slightly stronger (r=0.53) than between LB and MGs (r=0.38). No association between the diagnosed CM severity (mild vs severe) and fitness in mouse MGs was evident (Figure 4d).

**Figure 4.**
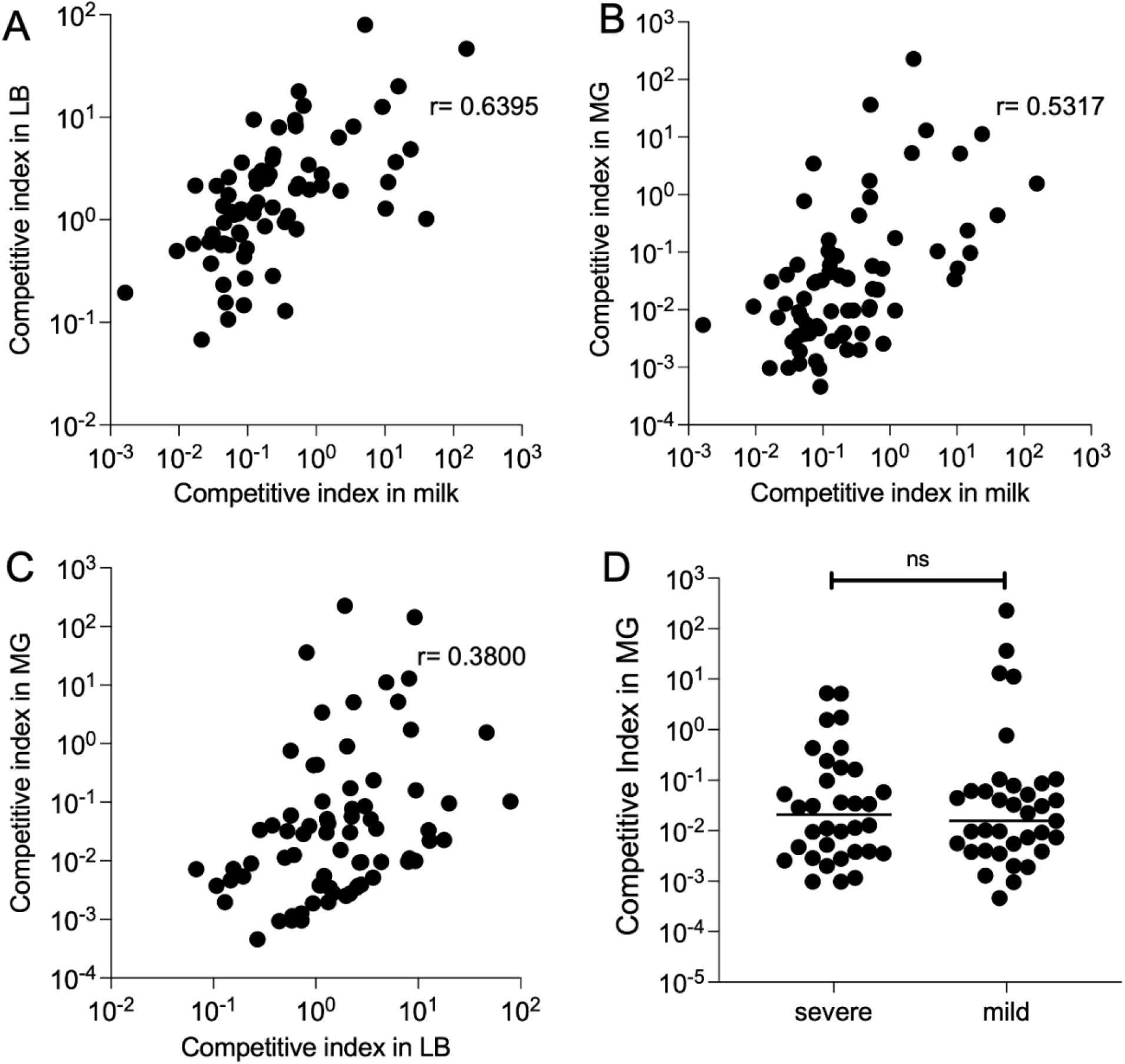
Competitive fitness of barcoded MAEC strains. All strains were inoculated together and grown in LB or unpasteurized cow’s milk (in duplicate) for 8 h, or in nine lactating MGs for 24 h. The bacteria were recovered, and their barcode plasmids were sequenced to determine their competitive indexes. (A) CI during *in vitro* growth in LB is strongly correlated with CI during growth in milk (*P*<0.0001 by Spearman rank correlation test). Each dot represents a single MAEC strain plotted at the mean competition index in both environments. (B) A slightly weaker positive correlation was detected between CI in milk and MGs (*P*<0.0001) and in (C) LB and MGs (*P* =0.0008) by Spearman rank correlation test. (D) CI in mouse MG infections for MAEC strains isolated from mild or severe CM cases. No significant difference was detected in these groups Student’s *t* test with Mann-Whitney correction (*P*=0.904).

Genes in the pan-genome that are associated with competitive fitness in milk and in mouse MGs were then identified. For this analysis, the top and bottom 30% of strains for each condition were separated based on their CI scores. Only five genes were positively associated with higher growth in milk, including a predicted inner membrane protein *yjeO,* three genes with unknown function, and an IS3 family transposase, *insK*. In mouse MGs, 38 genes were positively associated with increased competitive fitness (Supplementary Dataset 1). These included the bacterioferritin (*bfr*) and bacterioferritin-associated ferredoxin (*bfd*) genes that are involved in intracellular iron storage and mobilization (68). Notably, 14 of these genes are involved in an accessory type 2 secretion system found in two operons (*gspC-O* and *gspAB*) as well as the chitinase encoded by *chiA* (Table 4).

**Table 4.** Top genes* associated with MAEC fitness in mouse MG infections.

|  |  | Proportion of strains |  |  |
| --- | --- | --- | --- | --- |
|  |  | Most fit | Least fit | P-value |
| *rpiB* | Ribose-5-phosphate isomerase B | 0.812 | 0.313 | 3.85E-03 |
| *bfd* | Bacterioferritin-associated ferredoxin | 0.812 | 0.375 | 2.90E-02 |
| *bfr* | Bacterioferritin | 0.812 | 0.375 | 2.90E-02 |
| *gspB* | Putative general secretion pathway protein B | 0.812 | 0.375 | 2.90E-02 |
| *gspC* | Putative type II secretion system protein C | 0.812 | 0.375 | 2.90E-02 |
| *gspA* | Putative general secretion pathway protein A | 0.812 | 0.375 | 2.90E-02 |
| *gspF* | Putative type II secretion system protein F | 0.812 | 0.375 | 2.90E-02 |
| *gspG* | Putative type II secretion system protein G | 0.812 | 0.375 | 2.90E-02 |
| *gspD* | Putative secretin GspD | 0.812 | 0.375 | 2.90E-02 |
| *gspE* | Putative type II secretion system protein E | 0.812 | 0.375 | 2.90E-02 |
| *gspJ* | Putative type II secretion system protein J | 0.812 | 0.375 | 2.90E-02 |
| *gspH* | Putative type II secretion system protein H | 0.812 | 0.375 | 2.90E-02 |
| *gspI* | Putative type II secretion system protein I | 0.812 | 0.375 | 2.90E-02 |
| *gspO* | Type 4 prepilin-like proteins leader peptide-processing enzyme | 0.812 | 0.375 | 2.90E-02 |
| *gspL* | Putative type II secretion system protein L | 0.812 | 0.375 | 2.90E-02 |
| *gspM* | Putative type II secretion system protein M | 0.812 | 0.375 | 2.90E-02 |
| *gspK* | Putative type II secretion system protein K | 0.812 | 0.375 | 2.90E-02 |
| *chiA* | putative bifunctional chitinase/lysozyme | 0.812 | 0.375 | 2.90E-02 |
| *alsB* | D-allose-binding periplasmic protein | 0.812 | 0.375 | 2.90E-02 |

\*Excluding hypothetical proteins

As shown in the previous analysis (Table 3), this type 2 secretion system and *chiA* were also more frequently associated with pathogenic strains than with commensals. These associations suggested that ChiA might promote fitness of pathogenic strains. As ChiA proteins are known to enhance adherence of other bacteria to host cells or tissues, the role of ChiA in adhesion to cultured mammary alveolar epithelial (MAC-T) cells was investigated by creation of isogenic mutants. We chose to investigate three strains that were in our original study population that possess *chiA* (M45, M93, M111) as well as one additional strain that was not part of the original analysis (G1). M45 was among the top 30% most fit strains in MGs. M45 and M93 were isolated from mild cases of CM while M111 was isolated from a severe case. Strain G1 was isolated from a case of severe, gangrenous mastitis.

Wild type, *chiA* deletion mutants, or their complemented strains were added to MAC-T cells (MOI=10) for one hour and then the proportion of adherent cells measured. M45Δ*chiA* demonstrated an approximately 2-fold reduction compared to the wild-type parent (3.1×10^5^ vs 7.2×10^5^ CFUs, respectively) in attachment (Figure 5a). Attachment was restored upon reintroduction of *chiA* by plasmid complementation. A greater reduction was observed in M93 (>6-fold) between the wild type and mutant (Figure 5b), which was also able to be complemented. The slight reduction in attachment of M111Δ*chiA* compared to the wild-type strain was not statistically significant *P*=0.06) and complementation did not change adherence of this strain (Figure 5c). Deletion of *chiA* in G1 caused the largest decrease in attachment of all the MAEC strains we investigated. G1Δ*chiA* presented almost a 10-fold decrease when compared to the wild type as shown in Figure 5d. Wild-type levels of adherence were restored upon plasmid complementation in G1.

**Figure 5.**
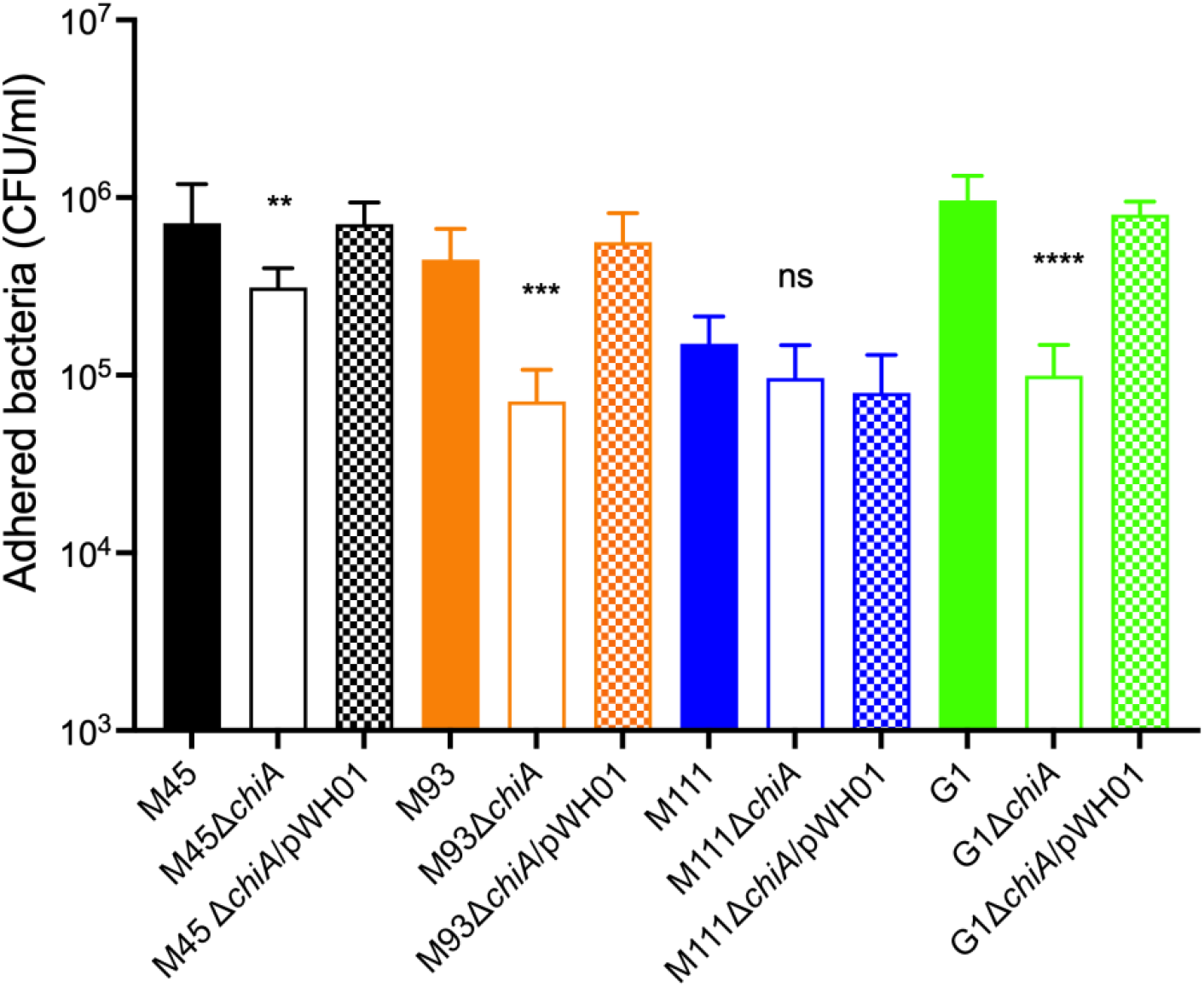
Role of ChiA in adherence of MAEC to mammary epithelial cells. Wild-type MAEC strains, their isogenic Δ*chiA* mutants, or the complemented mutant strains were tested for their ability to adhere to MAC-T cells at an MOI=10. One-way ANOVA with Tukey’s correction was used to determine significant differences between the wild-type and each Δ*chiA* mutant (\*\**P*=0.0082, *** *P*=0.0003, **** *P*<0.0001). These results are a representative experiment that was performed twice with six replicates per strain.

## Discussion

The features of pathogenic *E. coli* that differentiate them from non-pathogens remain incompletely understood, as is the relationship between bacterial fitness and the clinical disease that occurs during infection. Several recent studies have employed genome-wide association tools paired with clinical or experimental data to identify accessory genes associated with bacterial virulence, niche adaptation, AMR, and environmental persistence (69–75). In this study we sequenced 96 MAEC genomes, with the goal of identifying bacterial genes associated with differences in manifestation of bovine CM. We also compared genes associated with commensal vs pathogenic strains isolated from bovine and avian sources. We developed a barcoding system where multiple strains can be tracked as they compete in different conditions. This allowed us to quantify differences in fitness in specific controlled conditions and use these differences to identify genes associated with high and low-fitness strains. This approach enabled the identification of novel genes that may influence bacterial growth in multiple host environments. Our study differs from most bacterial GWAS in that it combines clinical and mouse infection assay data to identify a gene that was validated by functional studies.

Our data confirm the diversity of MAEC strains (Figure 1) and reveals the large pan-genome associated with these bacteria. From these accessory genes, we identified genes encoding adhesive and extracellular matrix structures associated with strains isolated from mastitis cases diagnosed as either mild or severe (Table 1). This includes the Yad fimbriae, which were positively associated with mastitis severity, and conversely, the PGA biofilm exopolysaccharide was negatively associated with mastitis severity. Yad fimbriae are also enriched among ExPEC strains of avian and human origin and are thought to promote adherence (76–81). Sustained inflammation in the MG is a main trigger for severe CM and bacterial factors that promote adherence would likely increase immune detection and signaling, leading to phagocytic cell recruitment. Alternatively, PGA may lessen CM severity by promoting biofilm formation, shielding inflammatory molecules on the surface of the bacteria and/or reducing invasion into epithelial cells.

MAEC are typically excluded from discussion of ExPEC strains more broadly. ExPEC exact an outsized disease burden in both humans and animals and are difficult to distinguish reliably from other *E. coli*. We did find several instances of MAEC strains possessing genes with demonstrated roles in ExPEC virulence (Figure 2). Although these virulence factors were not associated with CM severity generally, it does not rule out the possibility that they contribute to the virulence of individual strains in MGs, as we have previously demonstrated for the capsule and zinc uptake gene clusters of strain M12 (21, 30). Furthermore, additional factors continue to be recognized that contribute to ExPEC infections, and these may also be selected by the dairy environment and enriched in MAEC strains. The ferric dicitrate iron acquisition system is one such example (82). While their role in intramammary infection has been established, there is no previously known role in the pathogenesis of avian colibacillosis (Table 3). Our results suggest that the *fec* genes may also be enriched in avian-pathogenic strains, and they were recently demonstrated to enhance ExPEC urovirulence, suggesting that the *fec* system may be a factor in zoonotic spread of *E. coli* since it promotes fitness in multiple hosts and tissue types (82). The presence of these classic and newly appreciated ExPEC virulence genes in MAEC strains suggests that they occasionally infect humans and cause bloodstream or urinary tract infections.

We identified many different Inc plasmid groups in our MAEC strains and found that mild CM strains had significantly more Inc groups than severe CM strains (Figure 3). However, we did not investigate whether any virulence or AMR genes were carried on these plasmids as is frequently the case. The majority of the plasmids we detected belong to the IncF family which are usually conjugative (83), illustrating their potential to spread resistance in these populations. AMR is a significant health and environmental concern. However, only 12 strains carried one or more AMR genes. Two strains carried genes that confer resistance to six different classes of antimicrobials. Carriage of AMR genes did not appear to be associated with disease severity.

Unsurprisingly, our results indicate that those strains that are highly competitive *in vitro* tend to outcompete other strains *in vivo*, due to more rapid growth, direct antibacterial antagonism, or both (Figure 4). Interestingly, the correlation between competitive fitness in milk with MG infections than was slightly stronger than the correlation between fitness in LB and MG, suggesting that the ability to utilize nutrients or resist antimicrobial substances found in milk contributes to growth in lactating MGs. However, the lack of correlation we observed between bacterial fitness and CM severity (Figure 4d) illustrates that successful pathogens may replicate to high numbers without triggering deleterious responses in the host.

The *chiA* gene encoding a putative chitinase/chitin-binding protein was enriched in MAEC with higher fitness during mouse MG infections. This gene, along with the type II secretion system linked with it, was also associated with pathogenicity in the larger cohort of bovine and avian strains. However, they were not associated with CM severity. In the non-pathogenic K12 strain, transcription of this type II secretion system is normally repressed by the Hns protein, and the full *gsp* locus is needed for proper *chiA* secretion (84). ChiA has been implicated in the virulence of some adherent/invasive *E. coli* strains that cause colitis. This is not due to their chitinolytic activity, but rather because of chitin-binding domains that are found in the N-terminus. These domains mediate binding to chitinase-3-like-1 (CHI3L1), which is expressed on the surface of intestinal epithelial cells, leading to subsequent invasion of the bacteria (85, 86). Interestingly, expression of host chitin-like proteins is also induced by some bacterial pathogens (87). CHI3L1 regulates innate immune defenses against *Streptococcus pneumoniae* and *Pseudomonas aeruginosa* lung infections through inhibition of caspase-1-dependent macrophage pyroptosis (88, 89). Conversely, CHI3L1 expressed by intestinal epithelial cells during inflammatory bowel disease helps facilitate enteric bacterial infection.

Recently, CHI3L1 was also found in the milk secretions of quarters with bovine coliform mastitis (90). CHI3L1 gene expression is also increased in mouse MGs following *E. coli* infection. In knockout mice lacking CHI3L1, bacterial growth is not affected, but the influx of neutrophils into the lumen of the infected gland is reduced (90). It also promotes increased proliferation of mammary epithelial cells and reduces apoptosis (91). In the absence of CHI3L1, migration, maturation, and activation of macrophages is significantly impaired (92, 93). It seems likely that some MAEC strains may bind to CHI3L1 via ChiA, which could promote bacterial attachment and invasion and suppression of its inflammatory functions. In this way, bacterial fitness may be increased while limiting disease severity.

Alternatively, it is possible that ChiA contributes to infection through glycosidase activity, independently of its binding to host proteins. *Salmonella* chitinases modify glycans present in the extracellular matrix, uncovering mannose residues present on the epithelial surface and making them available for attachment through type I fimbriae (94, 95). They also increase survival inside phagocytes by dampening the expression of host antimicrobial responses in dendritic cells and macrophages (94). ChiA may play similar roles in MAEC colonization of MGs.

Our study has several limitations. First, the barcoding plasmid that we used to track mixed populations of bacteria may not behave identically in each strain’s unique genetic background. Although the plasmid has only one coding sequence for chloramphenicol resistance and is unlikely to affect gene expression broadly, the presence of the plasmid may interfere with stability of other native plasmids, which we have not examined. This may influence the fitness of these bacteria in unexpected ways. Secondly, competitive fitness of MAEC strains may be most relevant in natural environments in the presence of many other bacterial species other than *E. coli*. The ability to outcompete other *E. coli* strains may be less critical for MAEC than their ability to defend themselves against other diverse bacteria in the cattle GI tract, in soil, or ascending the MG teat canal. Future work should test what genes contribute to fitness in these environments and whether they can explain why some MAEC strains are more common, particularly those STs that we and others have identified. Finally, the results of our study may have been influenced by random factors such as when CM was diagnosed and scored. For example, a case that was detected early may have been diagnosed as mild whereas if diagnosed a few hours later may have been scored as moderate or severe. Diagnosis of CM is also inherently subjective, and the criteria may have been interpreted differently by individual farmers.

The wide range of severity and clinical presentation of MG infections by MAEC is impressive, and more studies directly characterizing the putative virulence factors of these strains are required. In this study, we have identified new accessory genes that could play a role in host specificity of these bacteria and influence disease outcomes. The role of ChiA in colonizing MGs as well as its functions in other ExPEC strains deserves further study, including its potential role in immune suppression and interaction with host structures. The ability of some MAEC strains to cross host species barriers and colonize different tissues underscores the importance of better understanding the diversity among this group of bacteria.

## Acknowledgements

We thank Paolo Moroni and Jennifer Wilson for providing MAEC isolates.

