## Supplementary Information for "Genes associated with fitness and disease severity in the pan-genome of mastitis-associated *Escherichia coli*"

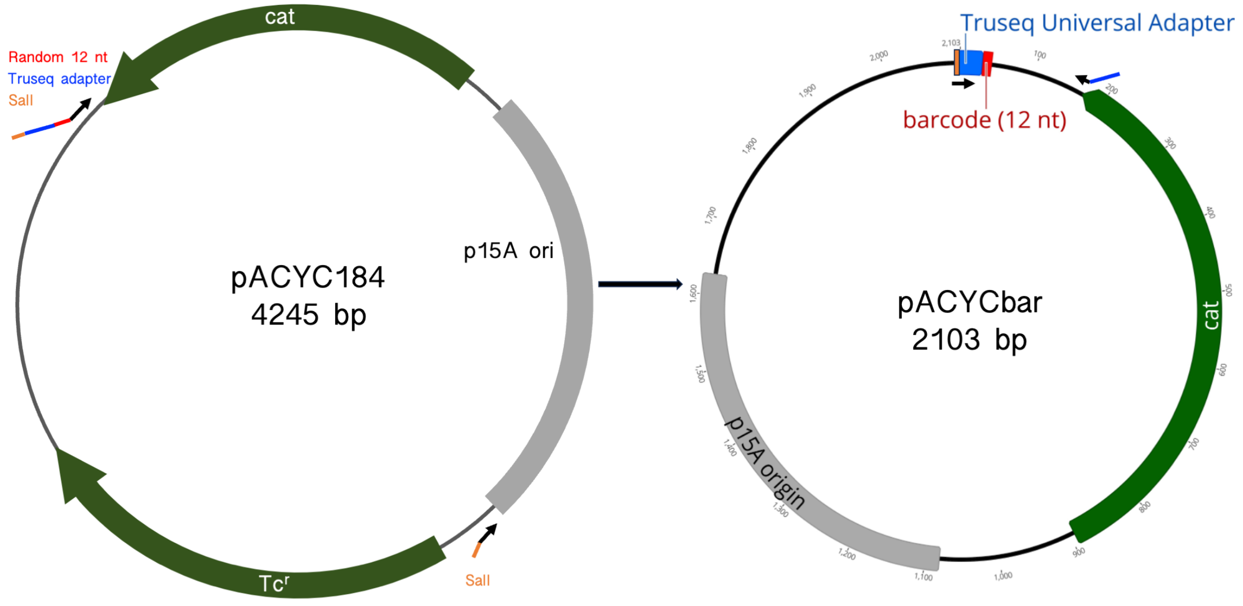


**Supplementary Figure 1. Development of barcode plasmid for multi-strain competition tests.** A fragment of pACYC184 was amplified by PCR to remove the tetracycline resistance gene. Barcodes were amplified from the resulting plasmids using primers that added the remainder of the Illumina adapter.

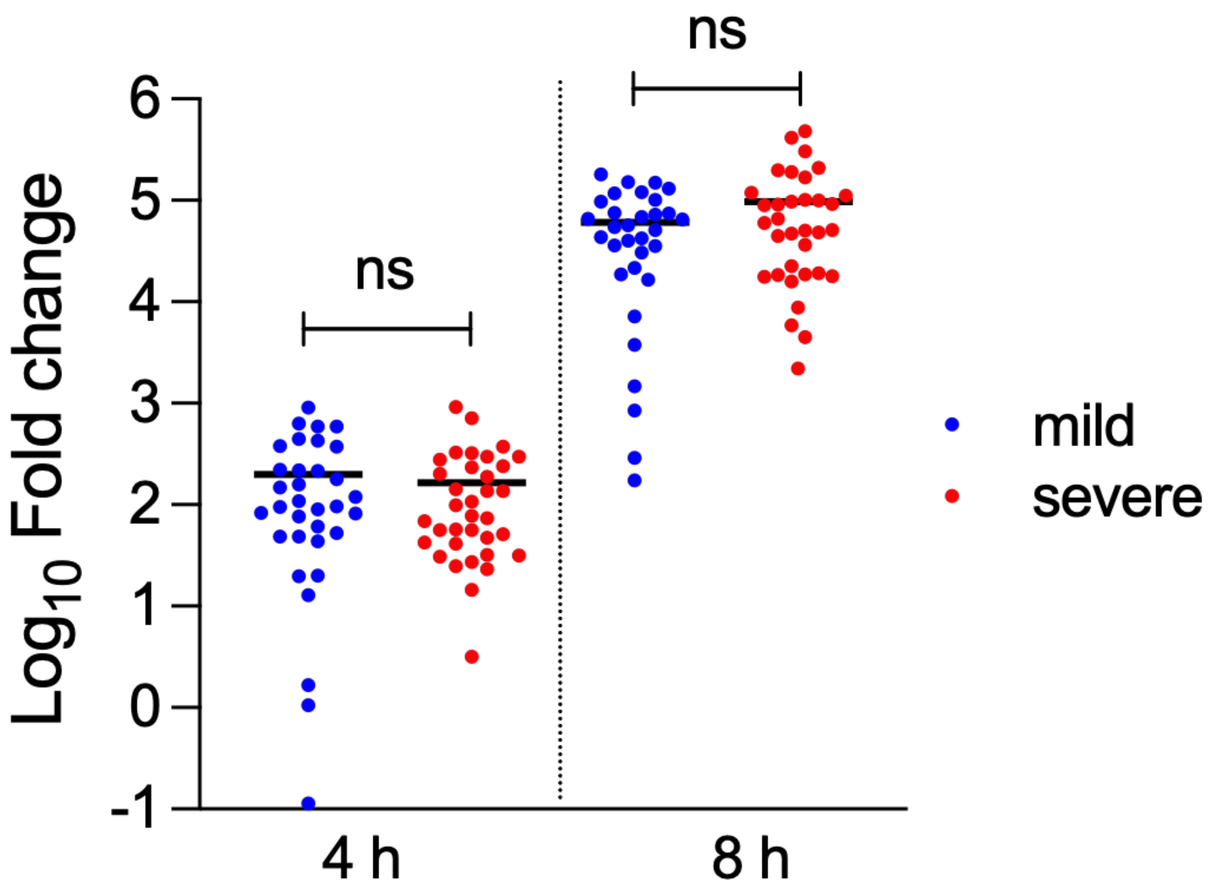


**Supplementary Figure 2. In vitro growth in milk of MAEC strains isolated from mild (n=34) or severe (n=32) mastitis.** Growth of each strain was measured in triplicate in whole unpasteurized milk at 4h and 8h. The fold change relative to the starting inoculum was determined by colony counts. There were no statistically significant differences between these two groups at either 4h or 8h by Student's *t* test with Mann-Whitney correction.

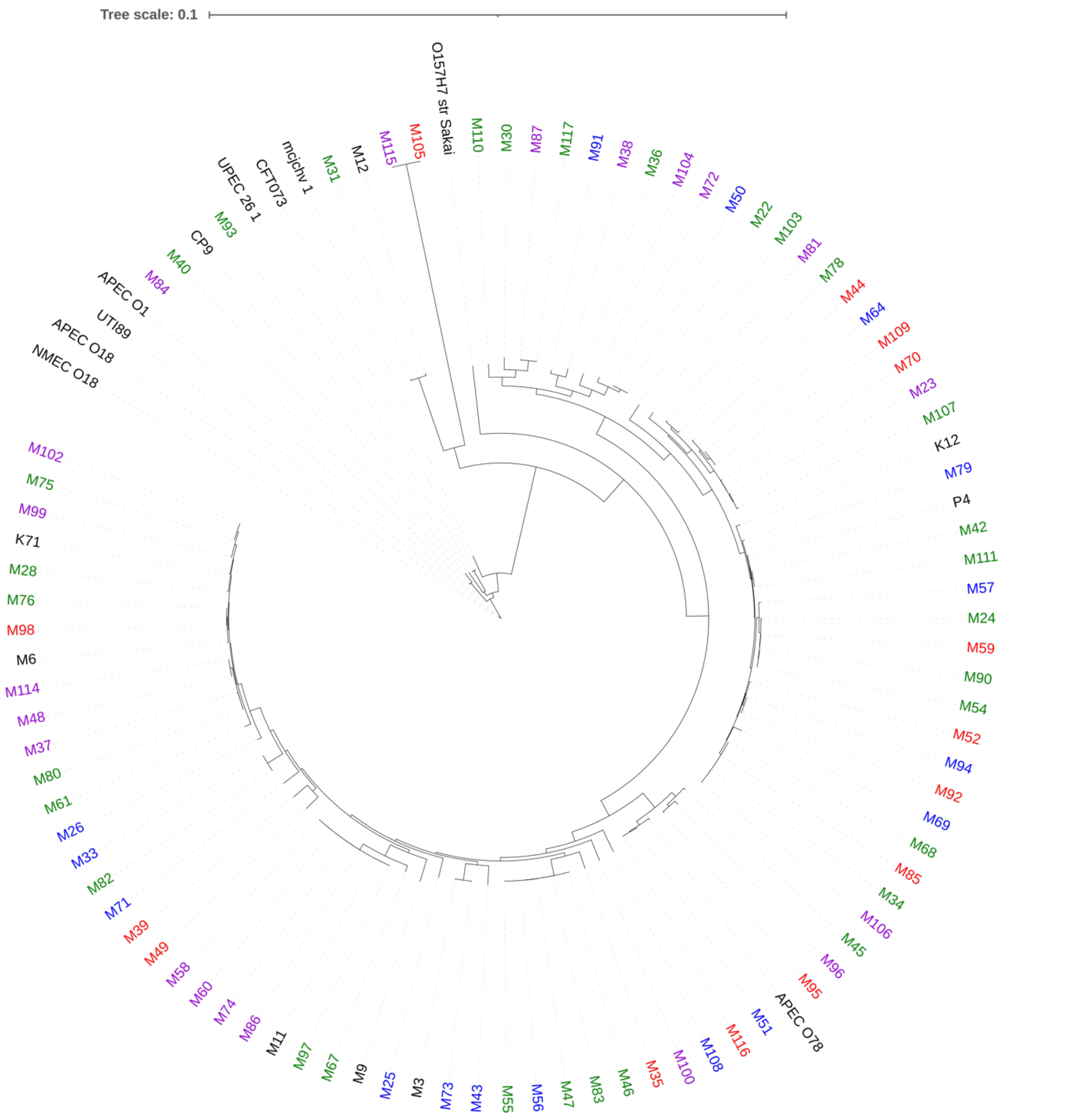


**Supplementary Figure 3. Core genome alignment does not reflect geographic location.** Strains isolated from Ontario, Quebec, Atlantic and Alberta and are labeled green, blue, purple, and red respectively. Strains labeled in black are reference strains.

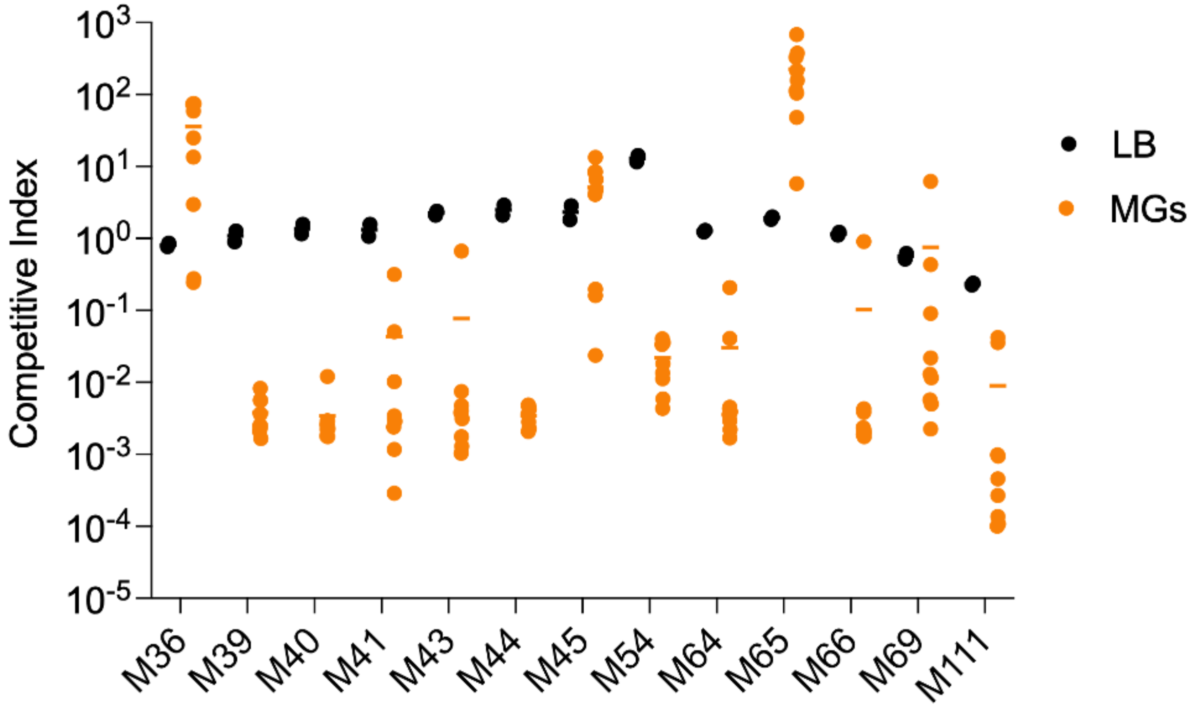


**Supplementary Figure 4. Competitive fitness of selected MAEC isolates.** A sample of strains is represented to show examples of highly competitive and less competitive strains. Data for individual MGs (n=9, orange circles) and replicate LB samples (n=2, black circles) are shown.

**Supplementary Table 1. Core and accessory genome statistics**

| Category | Frequency | Number |
|---|---|---|
| Core Genes | >98% | 3177 |
| Soft Core Genes | 95-98% | 263 |
| Shell Genes | 15-95% | 1523 |
| Cloud Genes | <15% | 10432 |
| Total Genes | 0-100% | 15395 |

**Supplementary Table 2. Antimicrobial resistance genes detected in MAEC genomes**

| Strain | A | B | C | M | P | T | Total |
|---|---|---|---|---|---|---|---|
| M79 | 3 | 1 | 2 | 1 | 1 | 1 | 9 |
| M96 | 2 | 1 | 3 | 0 | 2 | 1 | 9 |
| M51 | 4 | 0 | 2 | 0 | 1 | 1 | 8 |
| M102 | 3 | 1 | 2 | 0 | 0 | 1 | 7 |
| M110 | 4 | 1 | 2 | 0 | 0 | 0 | 7 |
| M75 | 3 | 1 | 2 | 0 | 0 | 0 | 6 |
| M99 | 2 | 1 | 2 | 0 | 0 | 1 | 6 |
| M39 | 2 | 0 | 1 | 0 | 0 | 1 | 4 |
| M70 | 1 | 0 | 2 | 0 | 0 | 0 | 3 |
| M97 | 2 | 0 | 0 | 0 | 0 | 0 | 2 |
| M87 | 0 | 0 | 0 | 0 | 0 | 1 | 1 |
| M92 | 0 | 0 | 0 | 0 | 1 | 0 | 1 |
| Total | 26 | 6 | 18 | 1 | 5 | 7 | |

Red=severe, Blue=mild CM
A=aminoglycoside, B=beta-lactam, C=antifolate, M=macrolide, P=phenicol, T=tetracycline
